# Motor training improves impaired cortico-cerebellar connectivity in cerebellar ataxia

**DOI:** 10.1101/2024.07.05.602300

**Authors:** Caroline Nettekoven, Rossitza Draganova, Katharina M Steiner, Sophia L Goericke, Andreas Deistung, Jürgen Konczak, Dagmar Timmann

**Author notes:** Correspondence to: Caroline Nettekoven Brain and Mind Institute, Western Interdisciplinary Research Building Western University London Ontario N6A 3K7, Canada.

## Abstract

People with cerebellar degeneration show characteristic ataxic motor impairments. Despite cerebellar dysfunction, they can still improve motor performance through sensorimotor training. Yet, how such training affects functional brain networks affected by cerebellar degeneration is unknown. We here investigated neuroplastic changes in the cortico-cerebellar network after a five-day forearm movement training in 40 patients with mild to severe cerebellar degeneration and 40 age- and sex-matched healthy controls. Participants were assigned to one of four motor training conditions, varying online visual feedback and explicit verbal feedback. Anatomical and resting-state fMRI was collected on the days before and after training. To overcome the limitations of standard brain templates that fail in the presence of severe anatomical abnormalities, we developed a specific template for comparing cerebellar patients with age-matched controls. Our new template reduced the spatial spread of cerebellar anatomical landmarks by 30% relative to existing templates and tripled fMRI noise classification accuracy. Using this pipeline, we found that patients showed impaired connectivity between cerebellar motor regions and neocortical visuomotor and premotor regions at baseline compared to controls, whereas their cortico-cortical connectivity remained intact. Training with vision strengthened connectivity in the cortico-cerebellar visuomotor network contralateral to the trained arm in all participants. Cerebellar patients exhibited additional increased connectivity ipsilateral to the training arm in this network. Further, training with explicit verbal feedback facilitated connectivity between a cerebellar cognitive region and dorsolateral prefrontal cortex. These results indicate that motor training in cerebellar degeneration leads to enhanced functional connectivity of the cortico-cerebellar network.

## Introduction

Dysfunction of the cerebellum causes deficits in the coordination and control of gait, posture, speech and fine motor movements. While the recovery of motor function after acute lesions of the cerebellum such as stroke typically happens within weeks and is often complete^1^, cerebellar neurodegeneration occurs over years and the ataxic motor symptoms worsen over time. For those affected by cerebellar degeneration, no medication has been proven effective and potential gene therapies are still in the early stages of development^2–4^. Existing therapies in patients with cerebellar degeneration therefore focus on mitigating symptoms through physical rehabilitation^4,5^. Continuous and intensive motor training has been shown to improve motor function in cerebellar patients^6–10^. However, no consensus exists on which type of training is most beneficial for cerebellar patients^4^. Understanding the neuroplastic changes that result from diverse types of motor training could help aid the development of tailored rehabilitation strategies for cerebellar patients.

Motor learning is considered an implicit learning process that relies on processing movement-relevant information from multiple sensory modalities such as vision and proprioception. It relies on an intact network comprising the cerebellum, the basal ganglia, sensorimotor cortex to modulate spinal or brain stem motor neurons. Given that the cerebellum is essential for motor learning and cerebellar degeneration impairs such learning, the utility of movement training programs has been questioned^11–13^. Indeed, empirical evidence shows that the ability to adapt to external forces, a form of implicit motor learning, can become abolished in the severe stages of cerebellar degeneration^14^. However, recognizing the degenerative process typically spans over years, motor training has also been shown to improve motor function in cerebellar patients at mild to moderate stages of cerebellar degeneration^6–10^. The extent of learning on a force adaptation task, has been found to correlate with the severity of ataxia in cerebellar patients with the most severely affected patients showing the smallest amount of learning^14^. To date, no physical rehabilitation programs exist that take the amount of residual sensorimotor function present in the patient into account.

It is well established that proprioceptive afferent signals are vital for motor learning and that the cerebellum receives large proprioceptive afferents through the spinocerebellar tracts^15^. In this context, it is noteworthy that proprioceptive perception is largely intact in people with cerebellar dysfunction as studies reported unimpaired limb position sense when the limb is passively rotated and no voluntary muscle activation is required^14,16^. However, cerebellar patients show impaired active position sense that relies on voluntary movement^16^. A training protocol that focuses on proprioceptive cues could potentially drive learning in cerebellar patients, but the extent to which patients can use proprioceptive information remains elusive.

Explicit error feedback given by a teacher, for example, can aid motor learning. This indicates that additional cognitive processes involving additional networks occur in parallel and can contribute to motor learning outcomes^17,18^. Research showed that cerebellar patients rely on explicit re-aiming strategies to counter rotations in a visuomotor adaptation task^19^, which implies that they may benefit from additional explicit feedback during motor training.

Moreover, animal work suggests that training can slow down the process of degeneration^20,21^. For people with cerebellar degeneration, motor training over several days led to structural changes in the neocortex, specifically grey matter increase in the premotor cortex^6,22^. Documenting changes in functional and structural connectivity can provide an additional, more detailed description of the initial impairment and associated training-related changes of the sensorimotor network. Functional activity and connectivity alterations have repeatedly been found in cerebellar patients^23–27^. Functional connectivity measured by resting-state fMRI has been shown to reliably dissociate patients from controls at 92% accuracy^28^. Functional connectivity changes might therefore hold potential for monitoring disease progression in response to putative therapeutic interventions.

However, a major obstacle to capturing connectivity changes in people with cerebellar degeneration are inadequate neuroimaging analysis pipelines. Normalizing degenerating brain structures to brain templates that are developed on neurologically healthy brains is challenging^29–32^, which limits our ability to detect patient-control differences at the group level. This is compounded by difficulties in aligning cerebellar structures^33^, low sensitivity of MRI acquisition coils to cerebellar signal^34^, and high noise levels induced by cardiac and respiratory cycles in areas close to the brain stem^35^. Hence, tailoring analysis pipelines to preserve cerebellar signal, particularly in patient populations, is crucial to revealing meaningful changes in connectivity.

In this study, we investigated the neuroplastic changes in cortico-cerebellar connectivity in a sample of patients with cerebellar degeneration induced by extensive motor training of their dominant right arm. The relevant behavioral and structural data have been published in Draganova et al., 2022^22^. The results showed that participants improved performance during visuomotor practice that was accompanied by an increase in premotor cortex grey matter.

However, a training focusing either on proprioceptive feedback, or explicit verbal feedback did not yield additional benefits. To delineate changes in functional connectivity associated with motor training, we developed a neuroimaging pipeline that substantially improved spatial localization by building a specialized template for comparing cerebellar patient data to data from healthy controls.

## Results

To understand the effect of different types of training on resting-state functional connectivity, we developed a five-day training protocol with varying task demands (see <u>4.2</u>). During training, 40 patients with cerebellar degeneration and 40 age-and sex-matched healthy controls performed elbow flexion movements in the horizontal plane with their right arm using a one degree of freedom single-joint manipulandum (described in^36^; Figure 1A). All participants were right-handed as assessed by the Edinburgh handedness scale^37^ and performed the task with their right arm. Participants were assigned pseudo-randomly to one of four training conditions, varying the type of feedback provided during and after the movement. During the movement, vision was either occluded through a mask (No Vision) or visual feedback was provided through a laser attached to the manipulandum (Vision). After the movement, the experimenter either provided explicit verbal feedback about the final position in relation to the target and how the target aim can be improved (Explicit Feedback) or no verbal feedback was provided (No Explicit Feedback). To quantify neural changes driven by training, structural and functional resting-state MRI data was acquired on the days before (pre) and after (post) the five-day training (Figure 1B). Behavioral and structural data have been reported in Draganova et al., 2022^22^.

**Figure 1.**
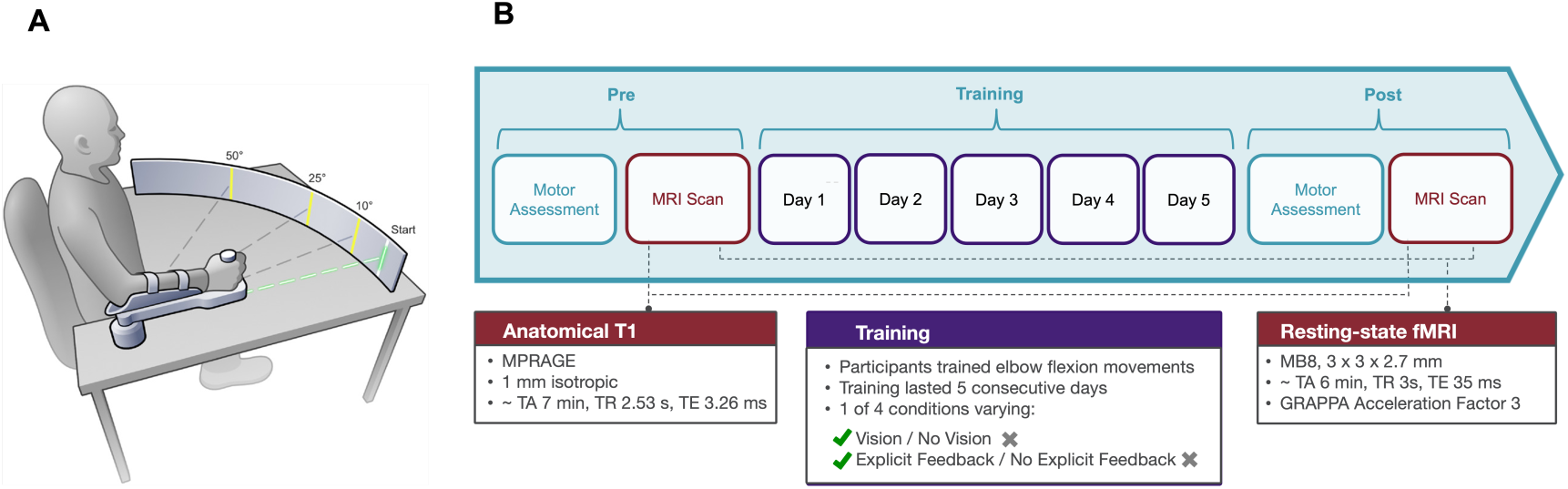
Study Design and Motor Training. **A.** Experimental setup of the single-joint manipulandum used for motor training. Start position was at 90elbow flexion. A low-intensity laser attached to the manipulandum provided visual feedback of arm position online. An optical encoder embedded in the housing recorded forearm position. Figure reproduced from Draganova et al., 2022. **B.** Participants participated in a five-day training with MRI sessions before (pre) and after training (post). Anatomical T1-weighted and resting-state fMRI data was collected during pre and post MRI sessions.

### Study-specific template improves spatial overlap

Anatomical abnormalities present in older adults and brains with cerebellar degeneration^30,38–44^, pose a challenge for accurate registrations to standard brain templates which are based on young, healthy adults. To improve spatial registration we therefore developed the **De**generation**Co**ntrol (DeCon) template, a template derived from the brains of cerebellar degeneration patients and controls in our study (Fig. 2A, see <u>4.4.1.1</u>). We used 40 anatomical scans, balanced across groups, timepoints and conditions (see Supplementary Table 1). To validate the use of the DeCon template, we examined the spatial overlap of anatomical landmarks after registration to the DeCon template (see <u>4.4.1.2</u>). We compared this to the spatial overlap after registration to two standard templates used widely in neuroimaging analysis of the cerebellum which are based on young, healthy brains: the MNI template^45^ and the SUIT template^33^. To ensure that our comparison is not biased towards the DeCon template, as it represents the average anatomy of those specific individuals, we repeated our analysis with anatomical scans from the 40 independent participants that were not used for template generation (validation sample).

**Figure 2.**
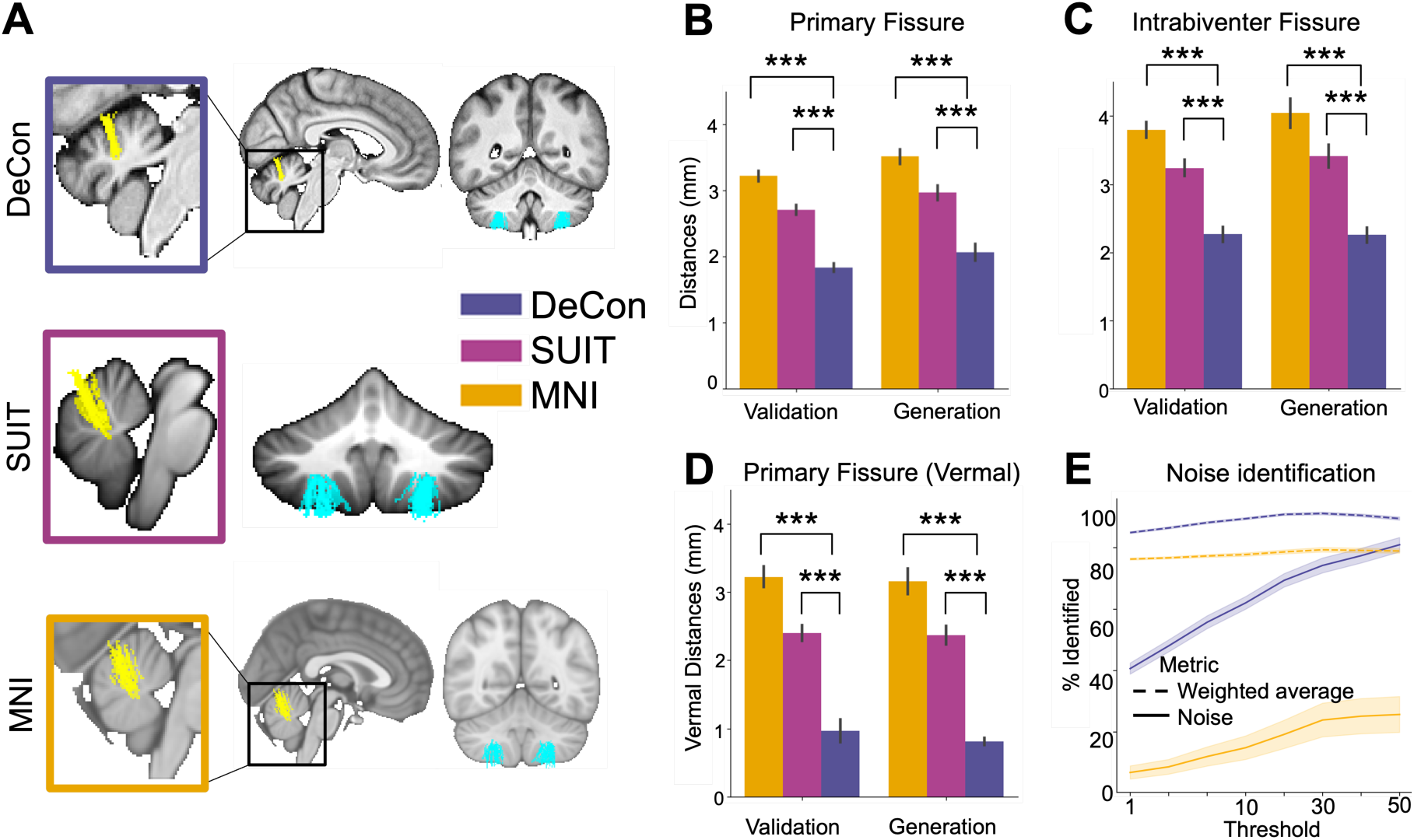
DeCon template improves anatomical alignment and functional localization. **A.** Primary (yellow) and intrabiventer (turquoise) fissures registered to the MNI, SUIT and DeCon template show the highest spatial overlap in DeCon space. Only fissures of participants that were not used for template generation are shown (n=40). **B-C.** Fissure overlap, computed as the average fissure distances between all possible pairs of participants, in MNI space (orange), SUIT space (magenta) and DeCon space (blue) is lowest in DeCon space. **B.** Primary fissure distance calculated within the whole cerebellum. **C.** Intrabiventer fissure distance calculated within the whole cerebellum. **D.** Primary fissure distance calculated within vermis. **E.** Percentage of correctly identified noise (solid line) and weighted ratio of correctly identified noise and signal (dashed line) when registering to the MNI template directly (orange) or via the DeCon template (blue). *** indicates significant p-values at 𝑝 < 0.001.

Two major cerebellar fissures, the primary and intrabiventer fissure were drawn on the native structural scan of each individual and transformed into DeCon, MNI and SUIT space. To quantify fissure overlap, we calculated the average fissure distances between all possible pairs of participants. The primary and intrabiventer fissures showed a 30% higher overlap in DeCon space than in both MNI and SUIT space (Fig. 2A). The largest reduction of fissure distances was found in the vermal portion of the primary fissure (see Supplementary Table 2). The primary fissure in the vermal portion was ∼0.9 mm apart in the DeCon template, compared to 2-3 mm in MNI and SUIT space (Fig. 2D). Paired t-tests demonstrated that the 70% reduction in fissure distances from MNI space and 60% reduction from SUIT space were significant for both the template generation sample (DeCon vs. MNI: 𝑡_39_= −15.49, 𝑝 = 2.97 × 10^−18^; DeCon vs. SUIT: 𝑡_39_ = −13.68, 𝑝 = 1.82 × 10^−16^) and the independent test sample (DeCon vs. MNI: 𝑡_39_ = −12.9, 𝑝 = 1.18 × 10^−15^; DeCon vs. SUIT: 𝑡_39_ = −7.88, 𝑝 = 1.36 × 10^−9^).

We next tested for a difference in alignment within the whole cerebellar volume. In the whole cerebellum, we found a smaller, but substantial decrease of 40% for MNI and 30% for SUIT space in primary fissure distance in the template sample (DeCon vs. MNI: 𝑡_39_= −14.91, 𝑝 = 1.06 × 10^−17^); DeCon vs. SUIT: 𝑡_39_= −13.43, 𝑝 = 3.28 × 10^−16^) and the validation sample (DeCon vs. MNI: 𝑡_39_ = −15.67, 𝑝 = 2.02 × 10^−18^; DeCon vs. SUIT: 𝑡_39_ = −11.32, 𝑝 = 6.83 × 10^−14^; Fig. 2B). This echoes observations from Schmahmann et al., 2000^46^, who noted that vermal sections of the primary fissure and other fissures are much more pronounced than hemispheric sections, resulting in blurrier fissure boundaries in cerebellar templates as the fissure extends more laterally^33^. In line with this, the DeCon template achieved a 40% and 30% reduction for intrabiventer fissure distance from the MNI template and the SUIT template, respectively (Generation sample, DeCon vs. MNI: 𝑡_39_ = −13.05, 𝑝 = 8.20 × 10^−16^; DeCon vs. SUIT: 𝑡_39_ = −10.11, 𝑝 = 1.90 × 10^−12^; Validation sample, DeCon vs. MNI: 𝑡_39_ =−18.46, 𝑝 = 7.08 × 10^−21^; DeCon vs. SUIT: 𝑡_39_ = −15.1, 𝑝 = 6.95 × 10^−18^).

To ensure that no systematic differences were present in anatomical alignment due to pathological anatomy, we tested for differences in fissure overlap between patients and controls in DeCon space using independent sample t-tests. We found no significant patient-control differences for the vermal primary fissure (𝑡_78_)_%_ = 1.05, 𝑝 = 0.30), the primary fissure across the whole cerebellum (𝑡_78_) = 1.09, 𝑝 = 0.28) or the intrabiventer fissures across the whole cerebellum (𝑡_78_) = 1.61, 𝑝 = 0.11). Our results demonstrate that the DeCon template substantially improves anatomical alignment in the cerebellum compared to existing templates, without favouring the anatomy of healthy control participants.

Finally, we asked whether the improvement in anatomical alignment results in better localization of functional data. For this, we used the automated classifier FIX^47^ to identify noise in the resting-state time series data (see <u>4.4.2.1</u>). Among several features used for noise classification, FIX uses masks of the major draining veins to identify features of vein-driven noise^47^. Because these masks are drawn in MNI-space and resampled to functional space, FIX’s ability to detect noise relies in part on an accurate mapping between MNI space and native functional space. We provided FIX with mappings between functional and MNI space estimated via the subject’s anatomical image only, or via the DeCon template <u>4.4.1.1</u> as an intermediate step between anatomical image and the MNI template. We then compared leave-one-out classification accuracy of FIX trained on hand-labelled signal and noise components across a range of classification thresholds in terms of correctly identified noise (true noise rate) and a weighted average between true noise rate and true signal rate (see <u>4.4.2.2</u>). Since FIX requires whole-brain registration, we only compared performance with the DeCon template, a whole-brain template. We did use the SUIT template as a comparison, since SUIT provides only a cerebellum template. Across classification thresholds, alignment via the DeCon template resulted in 46% more true noise components detected than alignment directly to MNI space (Paired t-test of true noise rate for DeCon vs. MNI 𝑡_40*_( = 10.51, 𝑝 = 4.54 × 10^−13^; Fig. 2E). Thus, the improved anatomical alignment to MNI space using the DeCon template yields higher accuracy in localizing functional data and hence, better noise feature detection. Finally, to remove structured noise from the resting-state data, we applied FIX-cleaning at threshold 30, where the weighted ratio of classification accuracy was highest (Fig. 2E).

### Cerebellar degeneration impairs connectivity with cerebellar regions

To understand the effects of cerebellar degeneration on the cortico-cerebellar motor network, we first examined functional connectivity at baseline. We quantified connectivity in a region-of-interest (ROI) approach as the Pearson’s correlation between the resting-state time courses of neocortical regions involved in motor control (primary motor cortex, M1; dorsal premotor cortex, PMd; posterior parietal cortex, PPC) and the contralateral cerebellar motor region (hand region M3) for both left and right (see <u>4.4.2.3</u>). We tested for significant effects of group (patient, control), regions (cortico-cortical, cortico-cerebellar, and cerebello-cerebellar) and a group x region interaction using a linear mixed effects model (see <u>4.5</u>). Patients showed significantly reduced connectivity at baseline (main effect of group: 𝐹_179,)"_ = 4.73, 𝑝 = 0.03; Fig. 3A) and this reduction depended on the region (group x region interaction: 𝐹_2,158_ = 162.55, 𝑝 = 4.52 × 10^−39^). Specifically, connectivity with cerebellar motor regions was lower in patients, while cortico-cortical connectivity remained intact (corrected two-sample t-test patients-control difference; cortico-cerebellar: 𝑡_77)_ = −3.07, 𝑝 = 0.02; cerebellar-cerebellar connectivity: 𝑡_77)_ = −2.73, 𝑝 = 0.02; cortico-cortical: 𝑡_77)_ = 0.51, 𝑝 = 1.00; Fig. 3B). Connectivity across all participants differed between regions (main effect of regions: 𝐹_2,158_ = 7.46, 𝑝 = 8.02 × 10^−4^), with strongest connectivity between the neocortical hemispheres, followed by cerebello-cerebellar connectivity and, finally, weakest connectivity between neocortex and cerebellum (all pairwise comparisons significant at 𝑡_156_ > 3.03, 𝑝 < 5.12 × 10^−3^).

**Figure 3.**
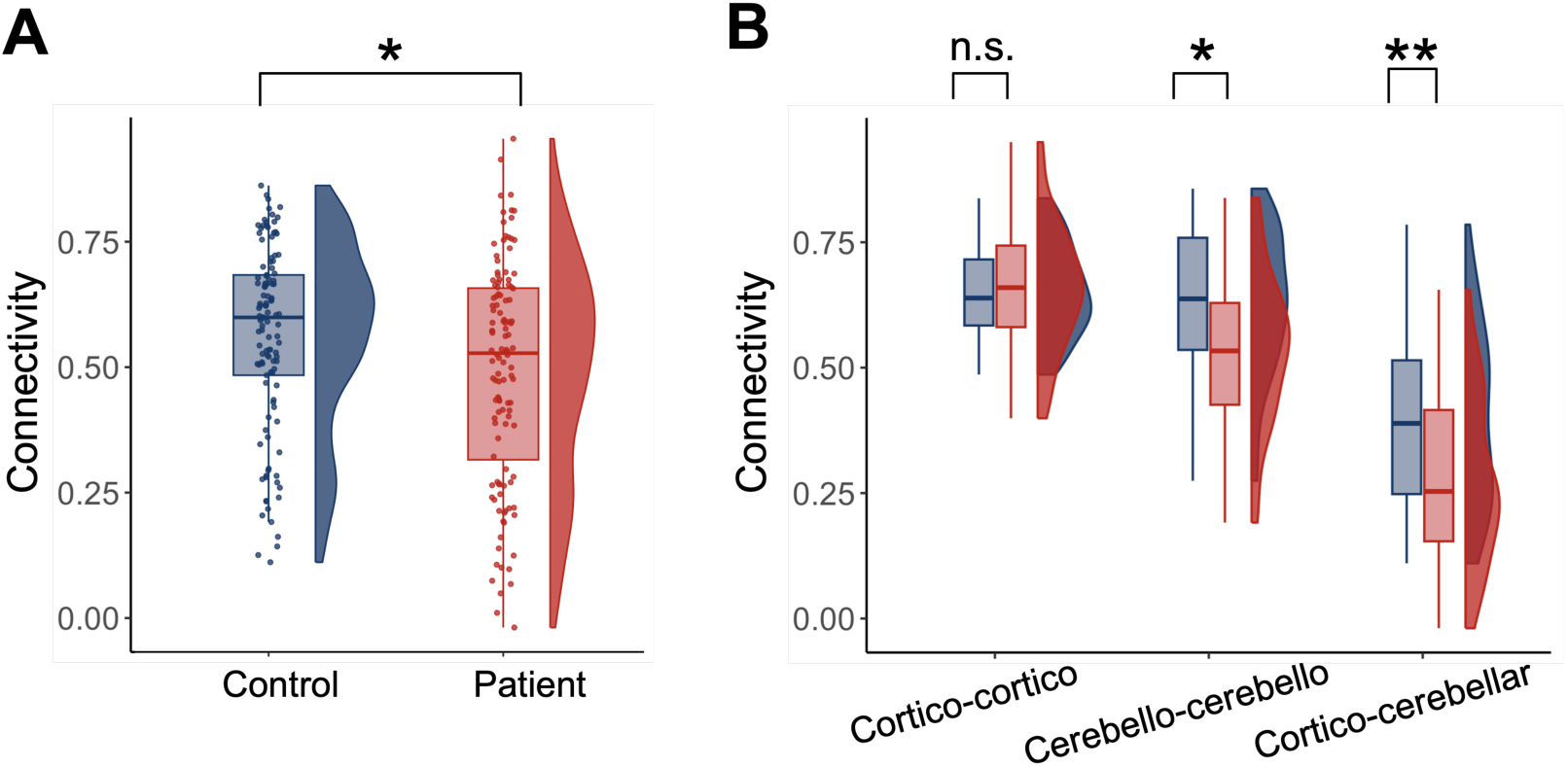
Impaired connectivity in the visuomotor network of patients. **A.** Connectivity of the visuomotor network differs at baseline between patients (red) and patients (blue). **B.** Connectivity shown separately for cortico-cortical, cerebello-cerebellar and cortico-cerebellar connectivity shows that connectivity reduction in patients is specific to connectivity with cerebellar regions. * indicates significant p-values at 𝑝 < 0.05. ** indicates significant p-values at 𝑝 < 0.01. Black asterisks indicate significant change in all participants.

### Training with visual feedback strengthens connectivity with bilateral premotor cortex in patients

Having established that cerebellar degeneration reduces connectivity in the cortico-cerebellar motor network, we then examined if prolonged training could reverse these impairments. We first examined training effects on connectivity with the dorsal premotor cortex (PMd), an area which has been shown to increase in grey matter through training in patients with cerebellar degeneration^6,22^. We modelled connectivity of PMd contralateral to the training hand with a cerebellar motor region ipsilateral to the training hand in a linear mixed effects model with fixed effects of group, session, vision (vision / no vision) and feedback (explicit feedback / no explicit feedback), and included all higher-order interaction terms, along with a random effect of subject. We observed a significant patient-control difference in connectivity (main effect group: 𝐹_1,78.76_ = 5.03, 𝑝 = 0.03; Fig. 4A). We also found a significant change in PMd- cerebellar connectivity (main effect session: 𝐹_1,78.15_ = 8.33, 𝑝 = 5.04 × 10^−3^), with higher connectivity after training in all participants (pre: 0.32 ± 0.21, post: 0.38 ± 0.23). We found that PMd-cerebellar connectivity changed depended on vision (session x vision interaction: 𝐹_1,78.15_ = 8.11, 𝑝 = 5.63 × 10^−3^), where participants who received visual feedback during training showed a significant increase in connectivity (vision: 𝑡_38_ = −3.86, 𝑝 = 8.50 × 10^−4^), whereas participants without visual feedback showed no significant change (no vision: 𝑡_38_ = −0.06, 𝑝 = 0.94). We observed no other significant main effects or higher-order interactions (all 𝐹 < 2.79, 𝑝 > 0.1).

**Figure 4.**
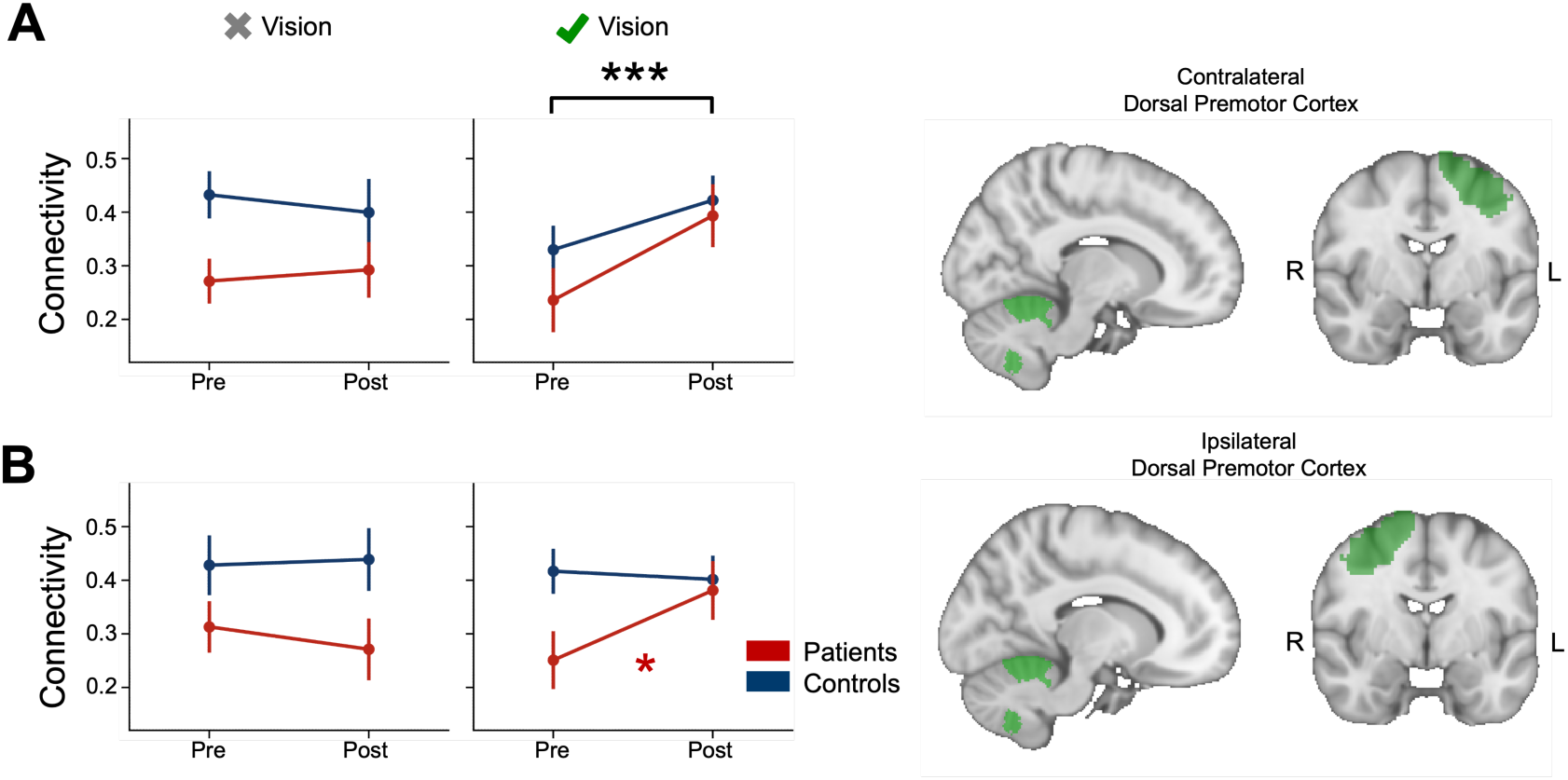
Connectivity change between dorsal premotor cortex (PMd) and cerebellar motor regions for training with and without vision. **A.** Left PMd connectivity with right cerebellar motor region. **B.** Right PMd connectivity with left cerebellar motor region. * indicates significant p-values at 𝑝 < 0.05. *** indicates significant p-values at 𝑝 < 0.001. Black *** indicates significant change in all participants, red * indicates significant change for cerebellar patient group only.

Previous investigations of training-related changes in PMd have reported grey matter increases ipsilateral to the training hand that were greater in patients than in controls^22^. We therefore assessed connectivity changes between right PMd and a left cerebellar motor region. We found that connectivity between right PMd and a left cerebellar motor region differed between groups (𝐹_1,78.82_ = 7.47, 𝑝 = 7.75 × 10^−3^, patients: 0.30 ± 0.23, controls: 0.40 ± 0.20, Fig. 4B), but there was no overall connectivity change (𝐹_1,78.20_ = 0.99, 𝑝 = 0.32). However, a significant interaction between group, session and vision indicated that connectivity change depended on group and visual feedback (group x session x vision interaction: 𝐹_1,78.20_= 5.746, 𝑝 = 0.01). Specifically, patients showed increases in connectivity when training with vision (𝑡_19_ = −3.18, 𝑝 = 0.02), but connectivity did not change in healthy controls training with vision (𝑡_18_ = −0.53, 𝑝 = 0.72), or in patients training without vision (𝑡_19_ = −0.37, 𝑝 = 0.72).

To test the specificity of our findings to cortico-cerebellar connectivity, we performed a control analysis on inter-hemispheric connectivity calculated between left PMd and right PMd. As expected, we found no patient-control difference (𝐹_1,78.97_ = 1.99, 𝑝 = 0.16), no significant change in connectivity (𝐹_1,78.67_ = 2.85, 𝑝 = 0.1) and no effect of vision on connectivity change (𝐹_1,78.97_ = 0.12, 𝑝 = 0.73). These control analyses confirmed that vision-dependent changes in PMd connectivity were specific to connectivity with cerebellar motor regions.

### Training with visual feedback increases cerebellar connectivity with contralateral posterior parietal cortex

In light of the key role the posterior parietal cortex plays in re-calibrating visually guided reaching movements^48^, we set out to test whether cerebellar connectivity with the posterior parietal cortex (PPC) increased when visual feedback was provided during training. We observed a significant difference in connectivity between patients and healthy control participants, with reduced connectivity in patients compared to controls (𝐹_1,78.96_ = 4.85, 𝑝 = 0.03; patients: 0.35 ± 0.21, controls: 0.44 ± 0.23; Fig. 5A). Connectivity between left PPC and right cerebellar motor region changed through training overall, with higher connectivity after training (𝐹_1,78.30_ = 4.95, 𝑝 = 0.03; pre: 0.37 ± 0.22, post: 0.42 ± 0.23). Connectivity also changed as a function of visual feedback (𝐹_1,78.30_ = 7.21, 𝑝 = 0.01). All participants who received visual feedback showed increased left PPC - right cerebellar connectivity after training (vision: 𝑡_38_ = −3.27, 𝑝 = 4.52 × 10^−3^; no vision: 𝑡_38_ = −0.31, 𝑝 = 0.76).

**Figure 5.**
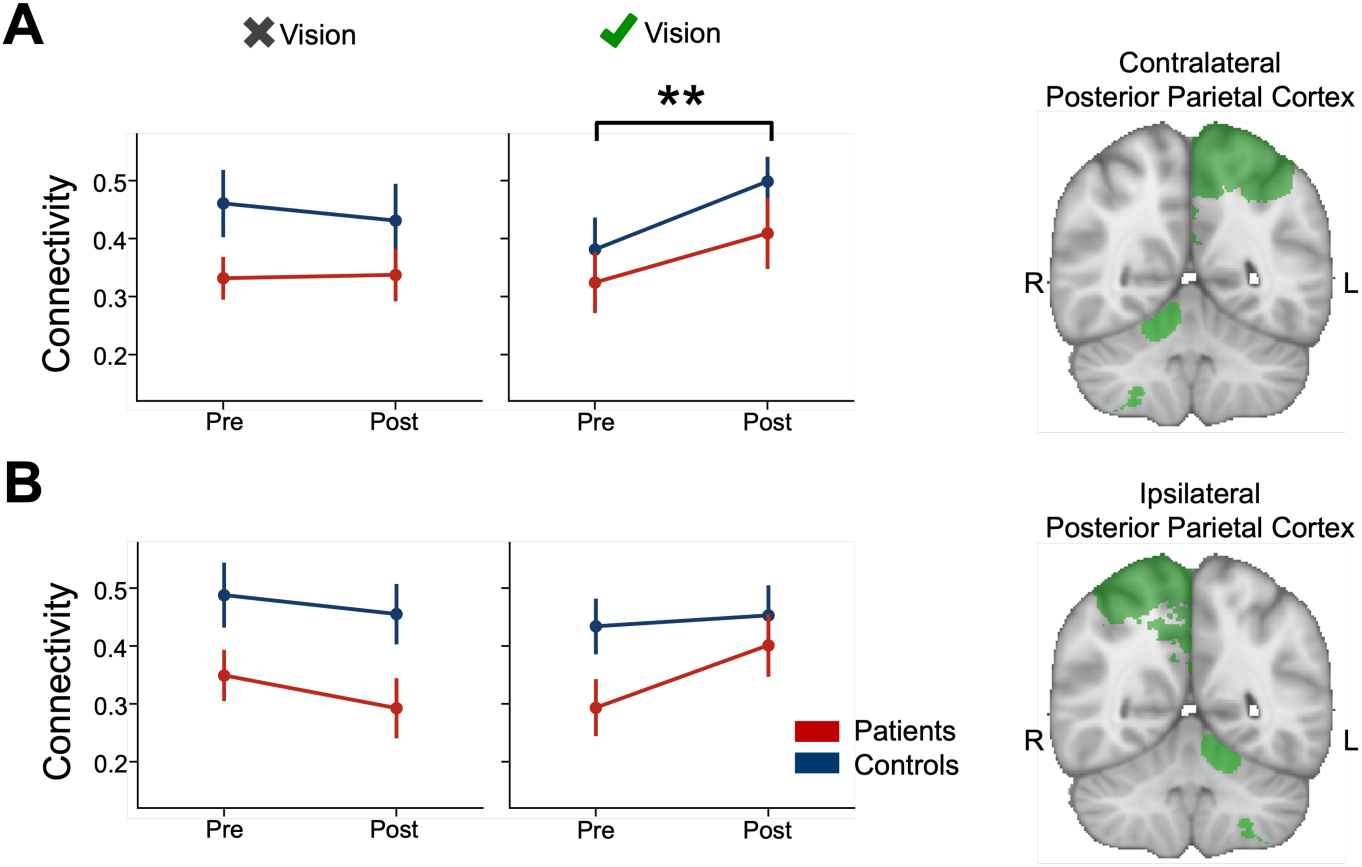
Connectivity change between posterior parietal cortex (PPC) and cerebellar motor regions when training with and without vision. **A.** Left PPC connectivity with cerebellar motor regions in the right hemisphere. **B.** Right PPC connectivity with cerebellar motor regions in the left hemisphere. * indicates significant p-values at 𝑝 < 0.05. ** indicates significant p-values at 𝑝 < 0.01.

Ipsilateral to the training hand, PPC connectivity with contralateral cerebellum showed a significant patient-control difference at baseline (𝐹_1,79.23_ = 9.11, 𝑝 = 3.42 × 10^−3^; Fig. 5B), but no connectivity change from baseline (𝐹_1,78.70_ = 0.26, 𝑝 = 0.61). Although a significant vision x session interaction indicated that vision affected connectivity change between these regions (𝐹_1,78.70_ = 5.94, 𝑝 = 0.02), post-hoc comparisons could not determine which condition drove this change (vision: 𝑡_38_ = 1.51, 𝑝 = 0.13, no vision: 𝑡_38_ = −1.82, 𝑝 = 0.13). A control analysis of left and right PPC connectivity found no patient-control differences (𝐹_1,79.34_ = 0.58, 𝑝 = 0.45), and no change in connectivity overall or as a function of visual feedback (𝐹_1,79.34_ = 1.28, 𝑝 = 0.26). Taken together, these results confirm the specificity of our findings to PPC-cerebellar connectivity.

### Motor training increases connectivity with contralateral primary motor cortex

Motor training has been shown to change connectivity between left M1 and right cerebellar cortex^49,50^. We tested whether practicing reaching movements, irrespective of training condition, changes functional connectivity between the right cerebellar and left M1 hand region. Indeed, we found that contralateral to the right hand, M1-cerebellar connectivity increased through training (𝐹_1,77.86_ = 4.64, 𝑝 = 0.03; Fig. 6A), such that connectivity was higher after training (pre: 0.22 ± 0.19; post: 0.27 ± 0.19). There was a significant patient-control difference in connectivity (𝐹_1,78.29_ = 8.69, 𝑝 = 4.21 × 10^−3^), but no difference in connectivity change between patients and controls (group x session interaction: 𝐹_1,77.86_ = 0.03, 𝑝 = 0.86). Previous studies indicated that M1-cerebellar connectivity change depended on learning^50^ but we did not find any studies that tested the influence of visual or explicit feedback. We found no effect of vision or explicit feedback on connectivity change (vision x session: 𝐹_1,77.86_ = 3.34, 𝑝 = 0.07, explicit feedback x session: 𝐹_1,77.86_ = 0.41, 𝑝 = 0.52).

**Figure 6.**
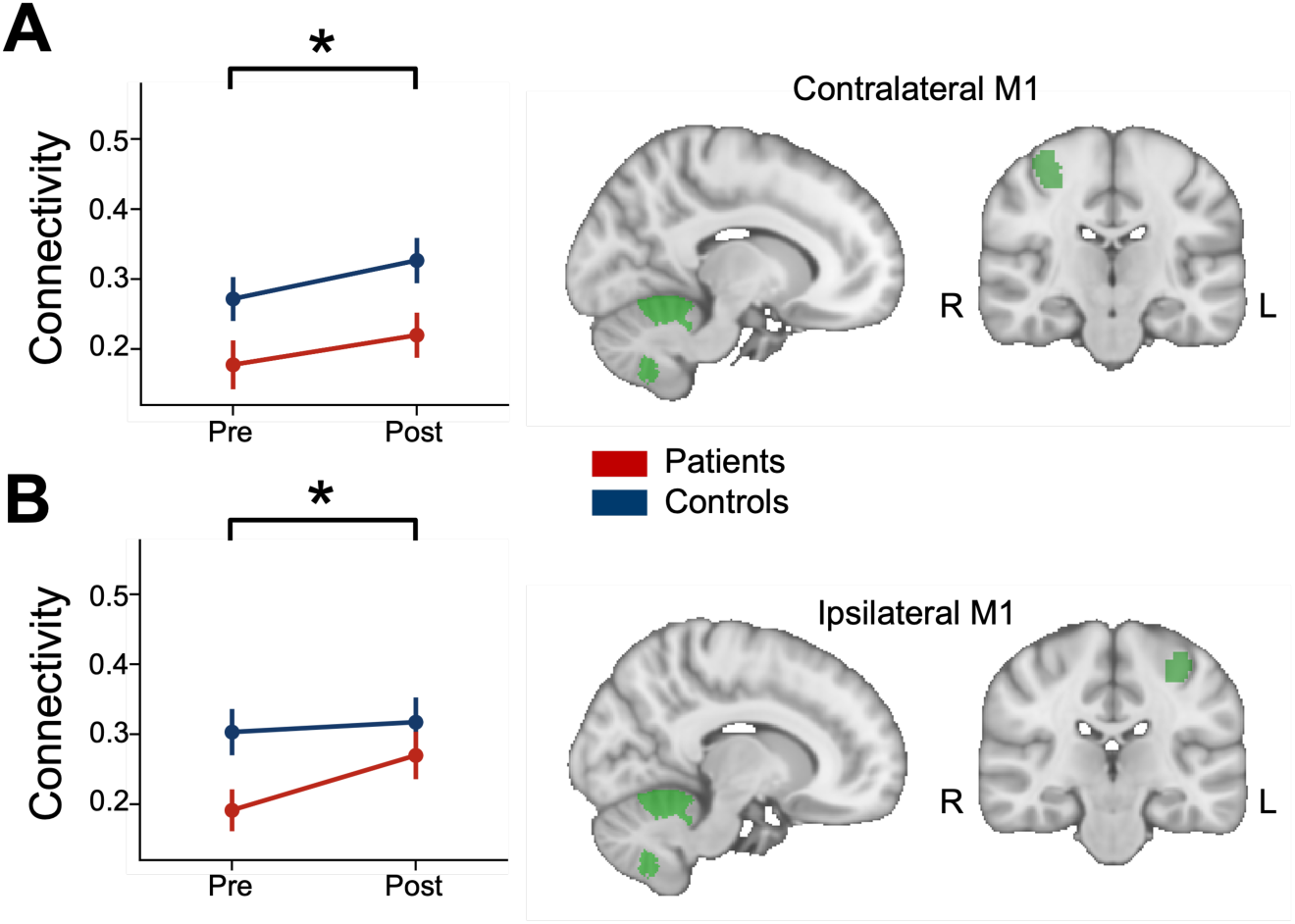
Connectivity change between M1 and cerebellar motor regions. **A.** Left M1 connectivity with right cerebellar motor region. **B.** Right M1 connectivity with left cerebellar motor region. * indicates significant p-values at 𝑝 < 0.05.

To test for the specificity of our findings to M1 contralateral to the training hand, we tested for connectivity changes from baseline of right M1 - left cerebellar connectivity. We found no significant connectivity change from baseline overall, suggesting that the training-related increase was specific to connectivity between left M1 and right cerebellar hand region (𝐹_1,78.46_ = 3.78, 𝑝 = 0.06; Fig. 6B). We also examined connectivity between the neocortical hemispheres for the M1 hand regions. Here, we observed no connectivity difference overall (𝐹_1,78.92_ = 2.91, 𝑝 = 0.09), but a significant increase in connectivity between left and right M1 (𝐹_1,78.63_ = 5.05, 𝑝 = 0.03).

Finally, we tested for a change in connectivity between left and a right cerebellar motor region through training or as a function of feedback. Our analysis showed a difference in connectivity between patients and controls at baseline (𝐹_1,78.97_ = 5.77, 𝑝 = 0.02). We also found a significant change in connectivity from baseline to post-training for all participants (𝐹_1,78.47_ = 8.13, 𝑝 = 5.571 × 10^−3^). This change depended on explicit feedback (𝐹_1,78.63_ = 5.40, 𝑝 = 0.02), but not on vision, group, or an interaction between these factors (all 𝐹 < 1.60, 𝑝 > 0.20). Thus, we cannot conclude that the overall change in M1-cerebellar connectivity was specifically driven by strengthening of cortico-cerebellar connectivity. However, the lack of significant higher-order interactions with vision suggest that our findings of PMd-cerebellar connectivity and PPC-cerebellar connectivity increases through training with vision are not driven by an increase in connectivity between cerebellar motor regions.

### Explicit feedback affects connectivity in cognitive cortico-cerebellar network

Cognitive regions of the neocortex have been implicated in strategy forming and explicit learning during visuomotor reaching tasks^51–53^. We therefore explored how explicit verbal feedback during training would affect connectivity between cognitive regions of the neocortex and the cerebellum. For this, we chose a region of interest from a novel functional atlas of the cerebellum, which was based on an aggregate of 7 extensive functional datasets^54^. We chose region D2 from the cerebellar atlas, which is a core multiple demand region that activates during the rapid processing of instructive signals in motor tasks^54^ and is located in lobules VI and VII along the superior posterior and the ansoparamedian fissures. We examined connectivity of D2 with the right dorsolateral prefrontal cortex (dlPFC), known to engage during strategy-based learning in adaptive reaching We observed a significant interaction between group, session and feedback (𝐹_1,78.83_ = 5.09, 𝑝 = 0.03; Fig. 7). Although paired t-tests of connectivity before and after training showed a significant decrease in connectivity for patients when training with explicit feedback (𝑡_19"_ = −2.14, 𝑝 = 0.04), but not without (𝑡_19_ = −1.05, 𝑝 = 0.31) these tests did not reach statistical significance when corrected for multiple comparisons (corrected p-values: 𝑝 = 0.18 in feedback conditions, 𝑝 = 0.42 in no feedback conditions). For connectivity between right dlPFC and left D2 or connectivity between left D2 and right D2, we found no significant interaction between group, session and feedback (right dlPFC - left cerebellar motor region: 𝐹_1,79.00_ = 3.95, 𝑝 = 0.05; right D2 - left D2: 𝐹_1,78.70_ = 0.08, 𝑝 = 0.78). Thus, any feedback-dependent effects on connectivity changes in patients and controls appear specific to right dlPFC connectivity with a right cerebellar cognitive region.

**Figure 7.**
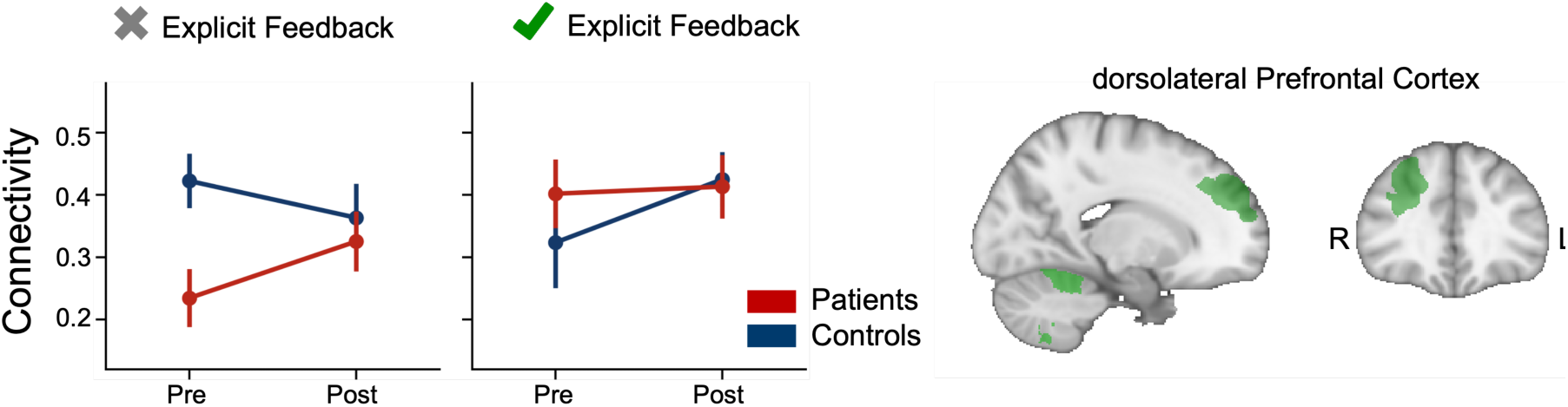
Connectivity change between right dorsolateral prefrontal cortex (dlPFC) and right cognitive cerebellar region (cerebellar atlas region D2 in Nettekoven et al., 2023^54^) training where explicit verbal feedback was given or not.

## Discussion

How cerebellar degeneration affects cortico-cerebellar connectivity and how therapeutic interventions can counter these effects are key clinical and neuroscientific questions that have yet to be addressed. In this study, we used a newly developed and carefully validated neuroimaging pipeline to demonstrate, for the first time, cortico-cerebellar connectivity alterations in cerebellar patients that ameliorate with motor training in a feedback-dependent manner. The main findings of the study are: First, at baseline, patients showed reduced connectivity of cerebellar motor regions with contralateral neocortical and cerebellar regions, while cortico-cortical connectivity was unimpaired. Second, visuomotor training led to increased connectivity between cerebellar motor and PMd contralaterally to the trained arm for all participants and ipsilaterally to the training arm for patients. Third, training with visual feedback increased connectivity with the PPC in all participants. These findings were highly specific to cortico-cerebellar connectivity. They were not driven by changes in neocortical connectivity, as left-right PMd connectivity and left-right PPC connectivity did not change overall.

To date, plastic changes after training interventions in cerebellar patients have only been demonstrated in regions not directly affected by degeneration^6^, including in a previous report of structural changes in this cohort^22^. These previous studies found grey matter increases in PMd, but were unable to demonstrate changes in cerebellar grey matter. Increased temporal synchrony of resting-state fluctuations in distant brain regions are thought to reflect increased communication of these regions^55,56^. This connectivity increase could be driven by strengthening of connections between brain regions involved in training, which remain detectable at rest post-training. In line with this, connectivity increases between task-relevant regions have been demonstrated in the visual cortex^57^, motor cortex^58^ and cerebellar cortex^59,60^. Given that the cerebellum projects to PMd^61^, our findings indicate that visuomotor training can take advantage of residual connections in the cortico-cerebellar network and induce lasting changes in the communication between the training-relevant regions.

The execution of visually guided movements involves the PPC^62^. In line with this, we found that visuomotor training increases functional connectivity between a cerebellar motor region and PPC contralateral to the trained hand. Strengthening of a network involving cerebellar regions and contralateral PPC has previously been reported in response to motor adaptation in healthy participants^63^. To our knowledge, our study is the first to demonstrate connectivity changes involving the PPC in cerebellar patients as well as healthy controls.

Previous research has found strengthening of connectivity between cerebellum and M1 contralateral to the training hand, when adaptive learning takes place, but not when no learning was induced^50^. This M1-cerebellar connectivity change, assessed through paired-pulse transcranial magnetic stimulation measures of cerebellar-brain inhibition, appears to rely on the presence of larger movement errors and does not manifest when introducing systematic errors gradually or randomly^49^. Here, we found M1-cerebellar resting-state connectivity increased regardless of task condition. These findings are in line with prior work, since all task conditions in this study tended to induce similar rates of learning, with differences between conditions appearing only in some target directions^22^. Moreover, the training conditions in this study were matched in all aspects of training protocol apart from visual and explicit verbal feedback, which have so far not been reported to induce differential changes in M1-cerebellar connectivity.

When exploring changes in a cortico-cerebellar network involved in higher-order cognitive aspects of motor training, we observed changes that differed between patients and controls and depended on the presence of explicit feedback. While these differences appear to be driven by a connectivity increase in the patient group when training without explicit feedback, post-hoc tests did not survive multiple comparisons. Although these findings might point to a potential malleability of cognitive cortico-cerebellar connectivity through targeted interventions, they will need to be replicated in an independent sample.

We were able to reveal highly specific connectivity changes by optimizing our analysis pipeline for maximising overlap of cerebellar structures and thereby preserving cerebellar signal. By creating a novel template for the analysis of cerebellar patient data, the *DeCon* template, we reduced spatial spread of anatomical landmarks in the cerebellum by 30-70% from existing templates, including the MNI template and the SUIT template, which had so far been the gold standard in cerebellar neuroimaging analysis^64^. When training a noise classifier on cerebellar resting-state fMRI data, mapping to MNI standard space via the DeCon template in a multi-stage deformation tripled classification accuracy. To enable other researchers to take advantage of this methodological advancement, we are making the template freely available under https://github.com/carobellum/DegenerationControlTemplate. We believe that it will improve anatomical and functional analysis of patient data and prove useful to the research and clinical community.

### Conclusions

Our results show that altered cortico-cerebellar connectivity can be ameliorated in cerebellar patients with training in a feedback-dependent manner. We used a newly developed and carefully validated brain template to reveal these subtle, but robust connectivity changes. These results demonstrate a fundamental step forward in understanding residual function in cerebellar degeneration patients and developing training protocols that take this knowledge into account. Further, the methodological advances presented here constitute a valuable addition to the tools available for cerebellar patient studies.

## Materials and methods

### Participants

41 patients with cerebellar degeneration and 44 neurologically healthy individuals participated in this study. One patient and two controls dropped out before study completion due to acute illness unrelated to the study, and two control participants were removed from analysis due to incidental findings, resulting in a final sample of 40 patients (age 55 ± 11.29 years, 21 females) and 40 age and sex-matched controls (age 55 ± 10.83 years, 20 females). For detailed study demographics, see Draganova et al., 2022^22^. Patients were diagnosed with a pure form of cerebellar cortical degeneration. Predominant diagnosis were spinocerebellar ataxia type 6, autosomal dominant cerebellar ataxia type 3, and sporadic adult-onset ataxia of unknown etiology. Ataxia severity was assessed using the clinical Scale for the Assessment and Rating of Ataxia^65^ (SARA). Participants were pseudo randomly assigned to one of four motor training conditions. Training groups were matched for sex, age and ataxia severity (SARA scores) and all participants were right-handed as assessed by the Edinburgh handedness scale^37^. The study was approved by the Ethics Committee of the Essen University Medical Center and all participants gave verbal and written informed consent prior to testing.

### Motor training

During a five-day motor training, participants performed elbow flexion movements in the horizontal plane using a single-joint manipulandum to targets arranged on a semicircular metal frame at 95 cm distance (Fig. 1A; manipulandum and task details can be found in^22,36^). Participants began each trial in a start position of 90-degree elbow flexion. After a start command, participants performed a single, ramp-like movement without online correction to the target. Participants held their arm in the end position for 4s before moving it back to the start position. Participants were instructed to perform swift and accurate movements. If movements were too slow or too fast according to previously calculated limits, the experimenter provided correctional feedback to reduce or increase movement speed.

On each training day, participants performed 100 trials to targets located counterclockwise at 10and 50, in blocks of 25 trials that alternated between the two targets. The target of the first block was balanced across participants of each training group and alternated each day. Participants began each training day by performing three familiarization trials to the first target of that day. After each block, participants rested for 3 minutes and after completing two blocks, participants relaxed their arm outside of the manipulandum.

To continuously drive performance, the width of the targets was reduced according to the movement accuracy achieved by the participant. Participants began at the lowest performance level with a target width of 4.5 cm, corresponding to 2.7on the frame. After performing five consecutive movements where the beam of the laser pointer was on target, the target width was reduced to the next smaller width (4.5 cm/2.7, 3.5 cm/2.11, 2.5 cm/1.5, 1.5 cm/0.9, 0.5 cm/0.3). Participants were instructed about the five levels prior to the experiment and notified by the experimenter as they advanced to the next level. Behavioural results have been reported previously^22,36^.

### Conditions

Participants were pseudo-randomly assigned to one of four motor training conditions, varying online visual feedback (Vision/No Vision) and post-movement verbal feedback (Expl. Feedback / No Expl. Feedback). 10 patients and 10 controls performed each condition (Vision | No Vision | Vision + Expl. Feedback | No Vision + Expl. Feedback). In conditions without vision, participants wore a mask that fully occluded vision. Here, the experimenter guided the arm of the participant from the start position to the target before each trial to enable the participant to memorize the target position. After moving back to the start position, participants performed the movement to the target. The experimenter informed participants verbally about the movement outcome, indicating whether the movement was “on target" or “target has not been reached”. In conditions with explicit feedback, participants received verbal feedback after each movement about the final joint position relative to the target, and strategies to minimize future movement errors (i.e., “Target was undershot by 5 degrees. Increase movement by 5 degrees”).

Each trial lasted 10s in the vision only condition and 15-20s when verbal feedback was given in addition to vision. In the conditions without vision, trials lasted 50-60s. Therefore, each training session with vision lasted 45-60 minutes, and 90-100 minutes without vision.

### MRI acquisition

Magnetic resonance imaging data was collected on the days before and after training (Fig. 1B) on a 3T combined MRI-PET scanner (Siemens Healthcare, Erlangen, Germany) with a 16- channel head coil with the same scanning protocol. Structural MRI data was acquired using a magnetization-prepared rapid acquisition gradient echo (MPRAGE) sequence with the following parameters: TR = 2530 ms, TE = 3.26 ms, inversion time = 1100 ms, flip angle = 7; FOV = 256 x 256 x 176mm^3^; bandwidth = 200 Hz/pixel; GRAPPA acceleration factor 2 and 48 reference lines; whole brain, acquisition time = 6:24 min. Resting-state fMRI data was acquired with the following parameters: TR/TE = 3000/35 ms, FOV = 282 x 282 x 134 mm^3^, bandwidth = 2312 Hz/pixel, voxel dimension = 3x3x2.7 mm^3^, GRAPPA acceleration factor 3 and 90 reference lines, whole brain, acquisition time = 5:17 min for a total of 100 volumes. During resting-state fMRI acquisition, participants fixated on a cross-hair image presented centrally on the screen. No functional data was acquired for one healthy control participant at both timepoints and for another healthy control participant at timepoint post due to scanner failure. The first subject was therefore excluded from functional analysis, while the second one was included for baseline connectivity but excluded for connectivity change analysis.

### MRI Analysis

All MRI data was processed using the Functional MRI of the Brain Software Library, (FSL; version 6.0.4;^66^). Structural images were additionally processed using advanced normalization tools (ANTs)^67^ and ITK-SNAP^68^.

### Structural MRI analysis

Structural images were reoriented to standard MNI orientation, cropped to remove superfluous neck tissue and corrected for RF/B1 field inhomogeneities using FSL’s fsl_anat^66^. Brain masks were generated using the Optimized Brain Extraction for Pathological Brains (optiBET^69^). To ensure precise tracing of the border between brain and non-brain tissue, in particular for degenerated brains, brain masks were corrected by hand by an experienced technician otherwise blinded to the study. Using the hand-corrected masks, the structural images were skull-stripped.

### Template Generation

Given the anatomical abnormalities present in brains with cerebellar degeneration and in scans of older adults^30,38–44^, it was important to use a registration template that is appropriate for patient brains and older adults. We therefore constructed a study-specific template from the scans of 40 participants. The other half of the sample was held out for validation purposes (see <u>4.4.1.2</u>). Scans that showed highly abnormal anatomy or particularly severe pathology, such as enlarged ventricles or strongly degenerated cerebellum, were not used for template generation, but included in the validation sample. To avoid biasing the template to a specific group or training condition, the template was constructed from twenty patient and twenty healthy control brains balanced across conditions (see Supplementary Table 1). Additionally, the scans were chosen to be representative of the mean sample age and sex distribution and balanced across scan time points.

The study-specific template was generated from the brain-extracted structural images using the iterative process implemented by the advanced normalization tools (ANTs, antsMultivariateTemplateConstruction2,^67^). As a starting point for the template construction, an existing group template derived from brains aged 55-59 was selected, which presented the closest match to the mean age of our sample (obtained from Neurodevelopmental MRI Database^70^). In brief, the structural images were rigidly, then linearly and finally nonlinearly registered to the starting template. The resampled images were averaged and deformed using the inverse average deformation to obtain a true mean shape image. This image served as the new starting template for the next iteration. To optimize convergence, ANTs downsamples and smoothes the images before registration and repeats the iterative process at progressively finer resolution and smoothing kernels. Our template underwent five refinement stages at 8x6x4x2x1 mm image resolution, 6x4x2x1x0 voxel smoothing kernels, and 500x200x100x70x20 maximum iterations. We hereby created a new template for comparing degenerated brains with control brains (DegenerationControl template, DeCon). To obtain a mapping between the DeCon template and MNI template, the DeCon template was nonlinearly registered to the MNI template using antsRegistrationSyn. The resulting nonlinear warp field was converted to an fsl-compatible format using the c3d_affine_tool from ITK-SNAP^68^ and FSL’s fslulils^66^.

All structural images were nonlinearly registered to the template using antsRegistrationSyn^67^. To enable comparisons between the DeCon template and existing templates, we additionally nonlinearly registered the structural images to the MNI template^45^ (nonlinear 6th generation MNI152 template or MNI152NLin6Asym available at https://www.templateflow.org/, hereafter referred to as ’MNI template’) and the SUIT template^33^ using antsRegistrationSyn^67^, all at 1mm resolution ( with precision type set to float and histogram matching turned on and all other parameters set to default values). Registration to the SUIT template was performed after isolating the cerebellum from the surrounding tissue in the structural scan using the SUIT isolate function^33^.

### Template Validation

We evaluated the degree of anatomical correspondence between cerebella in group space by calculating the spatial spread of anatomical landmarks of the cerebellum, using a procedure established by Diedrichsen, 2006^33^. As landmarks, we chose two major fissures, the primary fissure separating lobules V and VI and the intrabiventer fissure VIIIa and VIIIb. Fissures were drawn as surfaces on the structural scans in native space of each participant by an experienced technician otherwise blinded to the study. The fissures were then resampled to DeCon space, SUIT space and MNI space. Spatial overlap was calculated as the average distance between corresponding fissures of each possible pair of participants. Specifically, we computed the smallest distance of each point of fissure A to a point on the fissure B and averaged these first across all points of the fissure A, and then across all participants. Fissure alignment was calculated both for the entire cerebellum, using a cerebellar mask in group space, and restricted to the vermal section, using a mask obtained from the SUIT atlas^71^.

As a second evaluation criterion, we compared the ability of the automated noise classifier FIX^47^ to identify noise components of functional data using structural-to-MNI registrations estimated either directly to MNI space or with the DeCon template as an intermediate step. Since noise driven by vein fluctuations is dissociable from neural signal only by voxel location in major draining veins, FIX uses MNI-space masks of the major veins resampled to native functional space to identify vein noise^47^. Thus, FIX’s ability to detect noise relies in part on an accurate mapping between MNI space and native functional space. We therefore compared FIX’s classification accuracy using functional-to-MNI registrations estimated directly or via the DeCon template (see <u>4.4.2.2</u>).

### Functional MRI analysis

The resting-state functional MRI data acquired before and after the training days was preprocessed using steps largely matched to the UK Biobank processing pipeline^72^. The data was motion corrected using MCFLIRT^66^, slice-timing corrected using Fourier-space time series phase-shifting, stripped of non-brain voxels using BET (Brain Extraction Tool;^73^) and high-pass temporally filtered with a cut off frequency of 100s. No spatial smoothing was applied. Functional data were registered to the MNI space via the DeCon template and the structural scan of each subject and session. To this end, functional data were first linearly registered to the subject’s structural image acquired in the same session using boundary-based registration (implemented in FLIRT,^74–76^). The linear transformation was concatenated with the structural-to-DeCon nonlinear warp field and the DeCon-to-MNI nonlinear warp field into a functional-to-MNI space warp field.

### FIX Training

Structured noise was removed from functional data using a single-subject independent component analysis (ICA), implemented in MELODIC (Multivariate Exploratory Linear Optimized Decomposition into Independent Components) and the automated classifier FIX^47^. Because existing labelled training datasets differed substantially in their study population and acquisition parameters from our data, we created our own training set by manually classifying a subset of scans^47,77^. We selected 41 of the 157 scans in the dataset (26%), balanced across groups, conditions and timepoints for hand classification. To avoid optimizing our analysis pipeline to any set of scans, we minimized overlap between scanning sessions selected for template generation and for hand classification, with scans from eight subjects and timepoints included in both.

### FIX Evaluation

Classification accuracy was evaluated using leave-one-subject-out cross-validation where ICA components from a single hand-labelled subject are left out of the training process and the classification accuracy is then tested on the left-out subject^47^. Classification accuracy was summarized in terms of true positive rate ("TPR"), i.e. the percentage of correctly detected signal components, and true negative rate ("TNR"), i.e. the percentage of correctly detected noise components. A commonly used overall performance measure was calculated as the weighted average of TPR and TNR: (3 ∗ 𝑇𝑃𝑅 + 𝑇𝑁𝑅)/4^47,78,79^. Since FIX provides probabilistic signal and noise assignments and applies a threshold to obtain a binary assignment, classification accuracy was evaluated over a range of thresholds (threshold of 1, 2, 5, 10, 20, 30, 40 and 50). Since leave-one-out testing showed highest overall performance at threshold 30 (91.2 % accuracy), noise components identified at threshold 30 were removed from the functional data using FIX unique variance cleanup.

### Functional connectivity

We quantified functional connectivity between neocortex and cerebellum using a region of interest (ROI)-based approach for motor regions and regions involved in cognitive aspects of motor planning. For the neocortex, we derived an ROI of the left and right primary motor cortex (M1) hand area based on previous functional MRI data of hand movements^80^. A dorsal premotor cortex ROI (PMd) was defined for the left and right hemisphere from a tractography-based atlas^81^. A posterior parietal cortex (PPC) ROI was defined as left and right area 7 of the Juelich histological atlas^82^ thresholded at 15. Finally we defined an ROI for the right dorsolateral prefrontal cortex (dlPFC) as area 46d of the Sallet atlas^83^ thresholded at 25.

To define cerebellar ROIs, we used a novel functional atlas of the cerebellum based on multiple fMRI datasets that parcellates the cerebellum into 32 symmetrical functional regions^54^. We defined cerebellar motor ROIs as the left- and right-hand motor regions M3L and M3R, which activate strongly in response to movements of the ipsilateral hand. To explore connectivity changes in cerebellar cognitive regions in response to explicit verbal feedback, we used regions D2L and D2R. D2 was chosen because it shows highest activity during rapid processing of instructive signals in a motor task (see Supplementary Figure S2 in Nettekoven et al., 2023) and its specific involvement in working memory recall^84^, a cognitive function that deteriorates in cerebellar degeneration^85^. D2 also functionally connects to prefrontal regions of the neocortex^54^.

We quantified functional connectivity as the Pearson’s correlation coefficient between the resting-state time courses of two ROIs. To ensure normality, we used Fisher’s r-to-z-transformation on the connectivity values. For visualization purposes, we used non-normalized connectivity values.

### Statistical analysis

Mean ± SD is presented throughout. Statistical analyses were conducted using R (version 4.3.1^86^) and Python (version 3.9.10). We used paired t-tests to test for differences in fissure alignment and noise identification (scipy version 1.12.0^87^).

To test for connectivity differences at baseline, we constructed a linear mixed effects model of baseline connectivity values using the lme4 package (version 1.1-35.1). We averaged connectivity values within subject across regions, such that each subject had one cortico-cortical, one cortico-cerebellar and one cerebello-cerebellar connectivity value. We included fixed effects of group (patient, control), region (cortico-cortical, cortico-cerebellar, cerebello-cerebellar) and a group x region interaction term.

To investigate changes in functional connectivity and how these depend on group and training condition, we constructed separate linear mixed effects models for each pair of ROIs. We included timepoint, group, vision and feedback as fixed effects, along with their higher-order interaction terms. We allowed intercepts for different subjects to vary in the linear mixed effects models, to account for covarying residuals within subjects. Separate models of connectivity with M1, PMd, PPC and dlPFC were constructed, as the hypotheses regarding the specific fixed effects of interest were different for different regions. Residuals of all linear models were confirmed to be homoscedastic by visually inspecting the residuals plotted against the predicted values of each independent variable. Normality of fixed effects residuals and random effect residual was confirmed by visual inspection of the Q-Q plot using the ggpubr package (version 0.6.0). P-values were obtained with Type III tests calculated with Satterthwaite’s method (alpha = 0.05) implemented in the lmerTest package (version 3.1.3)^88^. In models with significant interactions, pairwise differences were assessed at each level using independent t-tests for patient-control differences and paired t-tests for comparing connectivity before and after training. P-values for post-hoc tests were corrected for multiple comparisons using the Benjamini–Hochberg method^89^, where all patient-control comparisons or all pre-post comparisons formed one family of tests. For testing connectivity differences at baseline, region combinations formed a family of tests. Since tests between regions are not independent, we corrected for multiple comparisons using the Benjamini-Yekutieli method^90^, as this method does not assume independence of tests.

## Data availability

The study-specific template is publicly available at https://github.com/carobellum/DegenerationControlTemplate. The template validation data is publicly available at https://github.com/carobellum/DegenerationConnectivity/tree/main/data along with the extracted timeseries data supporting the connectivity findings of this study. Due to data privacy restrictions, the raw MRI data has not been openly released, but can be made available upon request to the corresponding author.

## Code availability

The code for generating and validating the template, analyzing the functional connectivity data and for recreating the results and figures in this paper is publicly available as the GitHub repository https://github.com/carobellum/DegenerationConnectivity.

## Acknowledgements

We would like to acknowledge Jacob Levenstein for providing advice on template generation and Joern Diedrichsen for helpful discussions about template validation. Special thanks to Beate Brol for hand-correcting cerebellar masks and manually drawing cerebellar fissures.

## Funding

This study was funded by a research grant of the German Research Foundation awarded to J.1. K. and D. T. (DFG TI 239/14-1), and a research grant of the Bernd Fink Foundation awarded to J. K. and D. T..

## Author contributions

R.D., J.K. and D.T. designed the research; R.D., K.M.S., S.L.G. and A.D. performed the research; C.N. analyzed the data; C.N. wrote the first draft of the paper, C.N., R.D., J.K. and

D.T. revised the paper.

## Competing interests

The authors declare no competing interests.

---

**Supplementary Table 1.** Template generation and validation sample demographics. Demographic data for scans included into the study-specific template (generation sample) and held out from the template (validation sample). Generation scans were balanced across groups and conditions and approximately balanced across age, sex, and timepoint. Conditions are vision (V), no vision (NV), vision with explicit feedback (V+F), and no vision with explicit feedback (NV+F).

| Sample | Age |  | Sex |  | Group |  | Condition |  |  |  | Timepoint |  |
| --- | --- | --- | --- | --- | --- | --- | --- | --- | --- | --- | --- | --- |
|  | Mean | Std | M | F | Patient | Control | V | NV | V+F | NV+F | Pre | Post |
| Validation | 53.97 | 11.85 | 22 | 18 | 20 | 20 | 10 | 10 | 10 | 10 | 40 | 0 |
| Generation | 57.15 | 9.96 | 23 | 17 | 20 | 20 | 10 | 10 | 10 | 10 | 21 | 19 |

**Supplementary Table 2.** Fissure distances. Average pairwise fissure distances fur subjects included into the study-specific template (generation sample) and held out from the template (validation sample). Distances are given as mean (standard deviation).

| Sample | Fissure | Area | MNI template | SUIT template | DeCon template |
| --- | --- | --- | --- | --- | --- |
| Validation | Primary | Cerebellum | 3.80 (0.73) | 3.24 (0.74) | 2.26 (0.67) |
|  |  | Vermis | 3.22 (0.95) | 2.40 (0.74) | 0.96 (1.04) |
|  | Intrabiventer | Cerebellum | 3.22 (0.51) | 2.71 (0.48) | 1.83 (0.40) |
| Generation | Primary | Cerebellum | 4.04 (1.36) | 3.41 (1.05) | 2.25 (0.71) |
|  |  | Vermis | 3.16 (1.16) | 2.37 (0.85) | 0.81 (0.32) |
|  | Intrabiventer | Cerebellum | 3.51 (0.68) | 2.97 (0.70) | 2.06 (0.79) |

**Supplementary Table 3.**
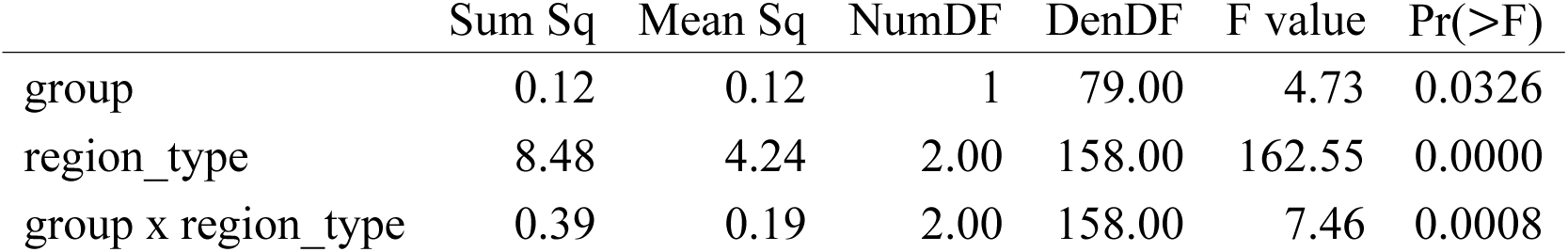
Linear mixed effects model for connectivity at baseline.

|  | Sum Sq | Mean Sq | NumDF | DenDF | F value | Pr(>F) |
| --- | --- | --- | --- | --- | --- | --- |
| group | 0.12 | 0.12 | 1 | 79.00 | 4.73 | 0.0326 |
| region_type | 8.48 | 4.24 | 2.00 | 158.00 | 162.55 | 0.0000 |
| group x region_type | 0.39 | 0.19 | 2.00 | 158.00 | 7.46 | 0.0008 |

**Supplementary Table 4.** Linear mixed effects model for connectivity between pmd-left and cereb-M3-right.

|  | Sum Sq | Mean Sq | NumDF | DenDF | F value | Pr(>F) |
| --- | --- | --- | --- | --- | --- | --- |
| group | 0.13 | 0.13 | 1 | 78.76 | 5.03 | 0.0277 |
| session | 0.21 | 0.21 | 1 | 78.15 | 8.33 | 0.0050 |
| vision | 0.00 | 0.00 | 1 | 78.76 | 0.04 | 0.8509 |
| feedback | 0.03 | 0.03 | 1 | 78.76 | 1.15 | 0.2872 |
| group x session | 0.05 | 0.05 | 1 | 78.15 | 1.99 | 0.1624 |
| group x vision | 0.03 | 0.03 | 1 | 78.76 | 1.37 | 0.2461 |
| session x vision | 0.21 | 0.21 | 1 | 78.15 | 8.11 | 0.0056 |
| group x feedback | 0.00 | 0.00 | 1 | 78.76 | 0.05 | 0.8223 |
| session x feedback | 0.00 | 0.00 | 1 | 78.15 | 0.11 | 0.7447 |
| vision x feedback | 0.00 | 0.00 | 1 | 78.76 | 0.19 | 0.6666 |
| group x session x vision | 0.00 | 0.00 | 1 | 78.15 | 0.12 | 0.7306 |
| group x session x feedback | 0.03 | 0.03 | 1 | 78.15 | 1.24 | 0.2681 |
| group x vision x feedback | 0.01 | 0.01 | 1 | 78.76 | 0.39 | 0.5346 |
| session x vision x feedback | 0.07 | 0.07 | 1 | 78.15 | 2.79 | 0.0986 |
| group x session x vision x feedback | 0.04 | 0.04 | 1 | 78.15 | 1.53 | 0.2199 |

**Supplementary Table 5.** Linear mixed effects model for connectivity between pmd-right and cereb-M3-left.

|  | Sum Sq | Mean Sq | NumDF | DenDF | F value | Pr(>F) |
| --- | --- | --- | --- | --- | --- | --- |
| group | 0.19 | 0.19 | 1 | 78.82 | 7.47 | 0.0078 |
| session | 0.03 | 0.03 | 1 | 78.20 | 0.99 | 0.3227 |
| vision | 0.00 | 0.00 | 1 | 78.82 | 0.08 | 0.7778 |
| feedback | 0.08 | 0.08 | 1 | 78.82 | 3.08 | 0.0831 |
| group x session | 0.04 | 0.04 | 1 | 78.20 | 1.63 | 0.2055 |
| group x vision | 0.02 | 0.02 | 1 | 78.82 | 0.79 | 0.3754 |
| session x vision | 0.06 | 0.06 | 1 | 78.20 | 2.34 | 0.1299 |
| group x feedback | 0.01 | 0.01 | 1 | 78.82 | 0.20 | 0.6546 |
| session x feedback | 0.07 | 0.07 | 1 | 78.20 | 2.61 | 0.1101 |
| vision x feedback | 0.00 | 0.00 | 1 | 78.82 | 0.00 | 0.9730 |
| group x session x vision | 0.15 | 0.15 | 1 | 78.20 | 5.75 | 0.0189 |
| group x session x feedback | 0.02 | 0.02 | 1 | 78.20 | 0.92 | 0.3398 |
| group x vision x feedback | 0.00 | 0.00 | 1 | 78.82 | 0.00 | 0.9911 |
| session x vision x feedback | 0.09 | 0.09 | 1 | 78.20 | 3.54 | 0.0637 |
| group x session x vision x feedback | 0.06 | 0.06 | 1 | 78.20 | 2.50 | 0.1178 |

**Supplementary Table 6.** Linear mixed effects model for connectivity between ppc-left and cereb- M3-right.

|  | Sum Sq | Mean Sq | NumDF | DenDF | F value | Pr(>F) |
| --- | --- | --- | --- | --- | --- | --- |
| group | 0.12 | 0.12 | 1 | 78.96 | 4.85 | 0.0305 |
| session | 0.12 | 0.12 | 1 | 78.30 | 4.95 | 0.0290 |
| vision | 0.00 | 0.00 | 1 | 78.96 | 0.06 | 0.8042 |
| group x session | 0.00 | 0.00 | 1 | 78.30 | 0.16 | 0.6935 |
| group x vision | 0.01 | 0.01 | 1 | 78.96 | 0.49 | 0.4867 |
| session x vision | 0.18 | 0.18 | 1 | 78.30 | 7.21 | 0.0088 |
| group x session x vision | 0.01 | 0.01 | 1 | 78.30 | 0.49 | 0.4874 |

**Supplementary Table 7.** Linear mixed effects model for connectivity between ppc-right and cereb-M3-left.

|  | Sum Sq | Mean Sq | NumDF | DenDF | F value | Pr(>F) |
| --- | --- | --- | --- | --- | --- | --- |
| group | 0.31 | 0.31 | 1 | 79.23 | 9.11 | 0.0034 |
| session | 0.01 | 0.01 | 1 | 78.70 | 0.26 | 0.6110 |
| vision | 0.00 | 0.00 | 1 | 79.23 | 0.01 | 0.9375 |
| group x session | 0.02 | 0.02 | 1 | 78.70 | 0.73 | 0.3965 |
| group x vision | 0.02 | 0.02 | 1 | 79.23 | 0.63 | 0.4304 |
| session x vision | 0.20 | 0.20 | 1 | 78.70 | 5.94 | 0.0171 |
| group x session x vision | 0.04 | 0.04 | 1 | 78.70 | 1.05 | 0.3092 |

**Supplementary Table8.** Linear mixed effects model for connectivity between m1-hand-left and cereb-M3-right.

|  | Sum Sq | Mean Sq | NumDF | DenDF | F value | Pr(>F) |
| --- | --- | --- | --- | --- | --- | --- |
| group | 0.15 | 0.15 | 1.00 | 78.29 | 8.69 | 0.0042 |
| session | 0.08 | 0.08 | 1.00 | 77.86 | 4.64 | 0.0343 |
| vision | 0.01 | 0.01 | 1.00 | 78.29 | 0.34 | 0.5608 |
| feedback | 0.04 | 0.04 | 1.00 | 78.29 | 2.11 | 0.1503 |
| group:session | 0.00 | 0.00 | 1.00 | 77.86 | 0.03 | 0.8624 |
| group:vision | 0.00 | 0.00 | 1.00 | 78.29 | 0.20 | 0.6572 |
| session:vision | 0.06 | 0.06 | 1.00 | 77.86 | 3.34 | 0.0716 |
| group:feedback | 0.04 | 0.04 | 1.00 | 78.29 | 2.09 | 0.1527 |
| session:feedback | 0.01 | 0.01 | 1.00 | 77.86 | 0.41 | 0.5254 |
| vision:feedback | 0.01 | 0.01 | 1.00 | 78.29 | 0.34 | 0.5590 |
| group:session:vision | 0.03 | 0.03 | 1.00 | 77.86 | 1.54 | 0.2181 |
| group:session:feedback | 0.01 | 0.01 | 1.00 | 77.86 | 0.77 | 0.3826 |
| group:vision:feedback | 0.00 | 0.00 | 1.00 | 78.29 | 0.14 | 0.7099 |
| session:vision:feedback | 0.00 | 0.00 | 1.00 | 77.86 | 0.14 | 0.7129 |
| group:session:vision:feedback | 0.00 | 0.00 | 1.00 | 77.86 | 0.00 | 0.9766 |

**Supplementary Table 9.**
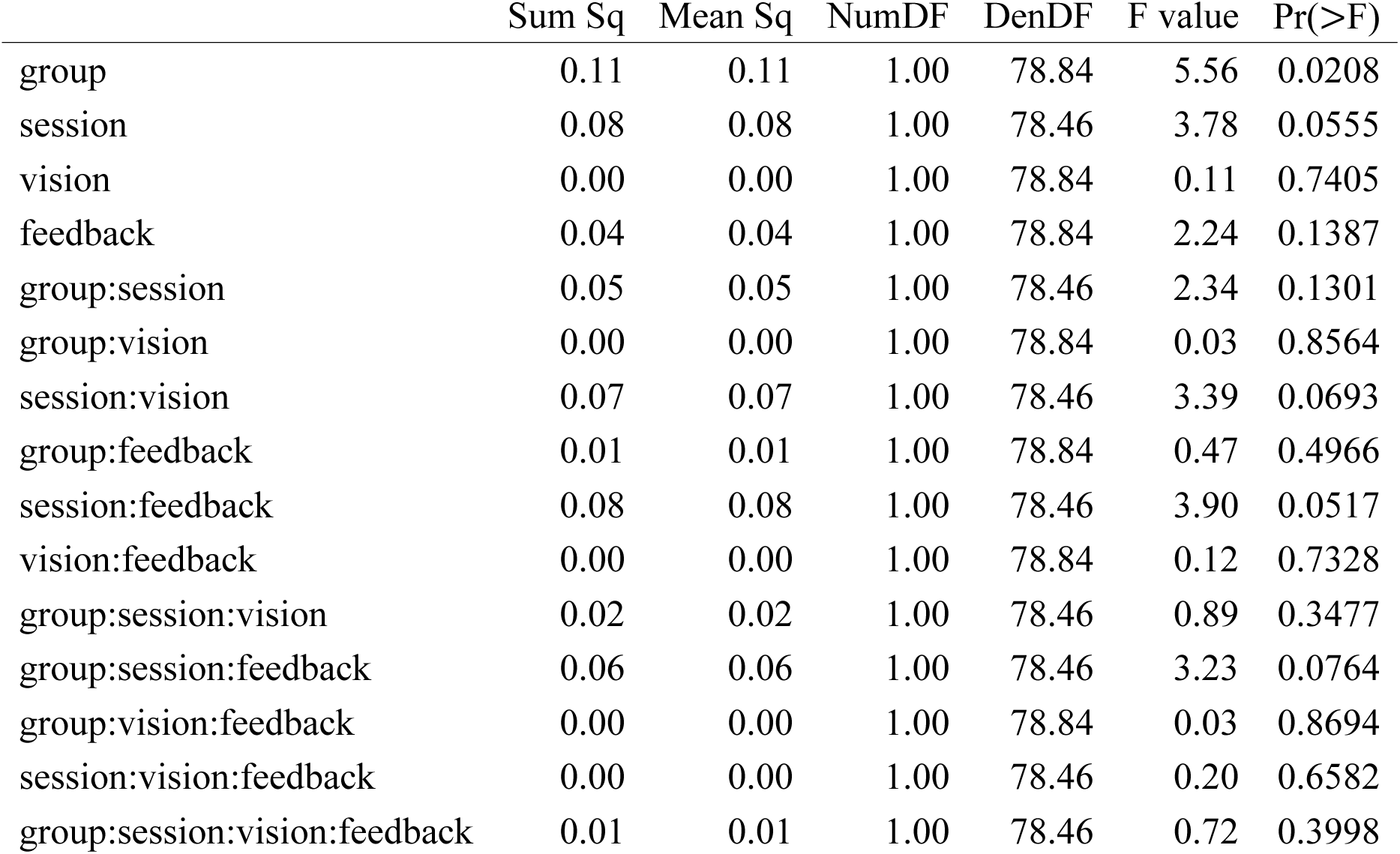
Linear mixed effects model for connectivity between m1-hand-right and cereb-M3-left.

**Supplementary Table 10.** Linear mixed effects model for connectivity between dlpfc-right and cereb-D2-right.

|  | Sum Sq | Mean Sq | NumDF | DenDF | F value | Pr(>F) |
| --- | --- | --- | --- | --- | --- | --- |
| group | 0.03 | 0.03 | 1 | 79.30 | 0.97 | 0.3286 |
| session | 0.06 | 0.06 | 1 | 78.83 | 1.69 | 0.1980 |
| feedback | 0.07 | 0.07 | 1 | 79.30 | 2.11 | 0.1503 |
| vision | 0.01 | 0.01 | 1 | 79.30 | 0.16 | 0.6868 |
| group x session | 0.01 | 0.01 | 1 | 78.83 | 0.25 | 0.6191 |
| group x feedback | 0.11 | 0.11 | 1 | 79.30 | 3.23 | 0.0759 |
| session x feedback | 0.01 | 0.01 | 1 | 78.83 | 0.34 | 0.5627 |
| group x vision | 0.00 | 0.00 | 1 | 79.30 | 0.00 | 0.9906 |
| session x vision | 0.03 | 0.03 | 1 | 78.83 | 0.87 | 0.3535 |
| feedback x vision | 0.01 | 0.01 | 1 | 79.30 | 0.21 | 0.6491 |
| group x session x feedback | 0.18 | 0.18 | 1 | 78.83 | 5.09 | 0.0269 |
| group x session x vision | 0.00 | 0.00 | 1 | 78.83 | 0.02 | 0.8943 |
| group x feedback x vision | 0.00 | 0.00 | 1 | 79.30 | 0.05 | 0.8245 |
| session x feedback x vision | 0.08 | 0.08 | 1 | 78.83 | 2.31 | 0.1322 |
| group x session x feedback x vision | 0.00 | 0.00 | 1 | 78.83 | 0.09 | 0.7601 |

## Notes

### Competing Interest Statement

The authors have declared no competing interest.

### Summary of Updates

PDF compiled incorrectly, so resubmitted PDF

